# Across-area synchronization supports feature integration in working memory

**DOI:** 10.1101/2021.06.09.447667

**Authors:** Joao Barbosa, Vahan Babushkin, Ainsley Temudo, Kartik K. Sreenivasan, Albert Compte

**Author notes:** Correspondence: Albert Compte.

## Abstract

Working memory function is severely limited. One key limitation that constrains the ability to maintain multiple items in working memory simultaneously is so-called swap errors. These errors occur when an inaccurate response is in fact accurate relative to a non-target stimulus, reflecting the failure to maintain the appropriate association or “binding” between the features that define one object (e.g., color and location). The mechanisms underlying feature binding in working memory remain unknown. Here, we tested the hypothesis that features are bound in memory through synchrony across feature-specific neural assemblies. We built a biophysical neural network model composed of two one-dimensional attractor networks – one for color and one for location – simulating feature storage in different cortical areas. Within each area, gamma oscillations were induced during bump attractor activity through the interplay of fast recurrent excitation and slower feedback inhibition. As a result, different memorized items were held at different phases of the network’s oscillation. These two areas were then reciprocally connected via weak cortico-cortical excitation, accomplishing binding between color and location through the synchronization of pairs of bumps across the two areas. Encoding and decoding of color-location associations was accomplished through rate coding, overcoming a long-standing limitation of binding through synchrony. In some simulations, swap errors arose: “color bumps” abruptly changed their phase relationship with “location bumps”. This model, which leverages the explanatory power of similar attractor models, specifies a plausible mechanism for feature binding and makes specific predictions about swap errors that are testable at behavioral and neurophysiological levels.

## Introduction

Working memory, our ability to hold information in mind for short time periods, is a hallmark of cognition but is severely limited on several fronts (Ma et al., 2014). Some of its limitations, such as its capacity, precision, or specific quantitative biases have been successfully accounted for by a family of biophysically-constrained models, mostly on the basis of a ring attractor network that maintains memoranda through sustained reverberatory neural activity (activity bumps) (Barbosa et al., 2020; Wimmer et al., 2014; Almeida et al., 2015; Qi et al., 2019; Nassar et al., 2018; Papadimitriou et al., 2015; Edin et al., 2009; Wei et al., 2012; Bouchacourt and Buschman, 2019; Compte et al., 2000). A feature of working memory that constrains the simultaneous storage of several items is the presence of swap errors (Schneegans and Bays, 2019). These errors occur when an inaccurate response to the target item is in fact accurate relative to a non-target item, reflecting the failure to maintain the appropriate association or “binding” between the separate features that define each item (e.g., color and location). The neural mechanisms supporting feature binding remain unclear, with different computational models implementing two alternative hypotheses (Matthey et al., 2015; Swan and Wyble, 2014; Pina et al., 2018; Schneegans et al., 2016; Schneegans and Bays, 2019; Raffone and Wolters, 2001).

The first type of models are based on selective synchronization (Raffone and Wolters, 2001; Pina et al., 2018). In these models, different neuronal populations selective to each feature that define an object are bound together through synchronized oscillatory activity. This would answer the longstanding question of how independently encoded features could be flexibly encoded as a single concept (Singer, 1999). Thanks to this flexibility, at least conceptually, these models do not suffer from combinatorial explosion as an increasing number of feature combinations are considered. There are however important questions about the biological plausibility of this hypothesis. Crucially, such a framework would need a temporal *encoder* that tags bound features by a “temporal code” and a temporal *decoder* that is able to distinguish which features are associated by detecting ensembles oscillating in precise synchrony. Both the encoder and decoder would thus depend on undefined biological mechanisms for spike coincidence detection (Shadlen and Movshon, 1999), which would struggle with the known high variability of neural spiking in sustained activity (Compte et al., 2003; Shafi et al., 2007). However, there is ample evidence for oscillatory dynamics during working memory. For instance, oscillatory activity in the gamma band (roughly defined between 30 hz – 100 Hz) increases during the mnemonic periods, both locally (Wimmer et al., 2016; Pesaran et al., 2002) and across sites (Lutzenberger et al., 2002; Kaiser et al., 2003; Kornblith et al., 2016; Palva et al., 2011), and further increases with memory load (Howard et al., 2003; van Vugt et al., 2010; Kornblith et al., 2016; Lundqvist et al., 2016). Importantly, gamma-band activity seems to play a functional role, as working memory binding performance is increased when transcranial stimulation at gamma frequency (40 hz) is applied at two different sites (left temporal and parietal), but only when in anti-phase (Tseng et al., 2016) in line with monkey electrophysiology showing that different items are stored in different oscillatory phases (Siegel et al., 2009) and the more general framework of phase-coding in working memory (Fell and Axmacher, 2011).

Another class of models achieve feature binding through “conjunction neurons” - neurons that are selective to all features being bound. Since neurons with mixed selectivity are ubiquitous in the brain (Fusi et al., 2016; Rigotti et al., 2013), these models seem more biologically plausible than those relying on unrealistically precise spike synchronization. Nevertheless, they suffer from some important limitations. First, the number of possible combinations explode quickly with an increasing number of features (Matthey et al., 2015; Schneegans et al., 2016; Schneegans and Bays, 2017, 2019) (but see (Swan and Wyble, 2014)). Second, these models do not have independent storage systems for each feature that define an object, to which there is converging evidence (Delvenne and Bruyer, 2004; Olson and Jiang, 2002; Parra et al., 2011; Xu, 2002; Wheeler and Treisman, 2002; Fougnie and Alvarez, 2011; Bays et al., 2011b). See (Schneegans and Bays, 2019; Ma et al., 2014) for recent reviews on the experimental evidence that should constrain multi-item working memory models, in particular those aiming to explain feature binding.

Here, we propose a hybrid model that overcomes several limitations from both types of models. We connected two ring attractor networks – one ring representing and memorizing colors and another ring storing locations – via weak excitation. This is an explicit implementation of the independent storage of individual features, where each feature might be represented in different cortical areas (e.g., color in inferior temporal cortex and location in posterior parietal cortex). Within each area, oscillatory mnemonic activity occured naturally through the interplay between fast recurrent excitation and slower inhibitory feedback. Feature binding was accomplished through the selective synchronization of pairs of bumps across the two networks. Furthermore, encoding and decoding of specific color-location associations was accomplished through rate coding. Our hybrid model of rate/temporal coding shares the rich explanatory power of classical ring-attractor models of working memory (Barbosa et al., 2020; Wimmer et al., 2014; Almeida et al., 2015; Qi et al., 2019; Nassar et al., 2018; Papadimitriou et al., 2015; Edin et al., 2009; Wei et al., 2012; Bouchacourt and Buschman, 2019) and derives new predictions that can be tested on multiple levels.

## Materials and Methods

### Neural Network Model

We extended a previously proposed computational model (Compte et al., 2000). In particular, we connected two one-dimensional ring networks via weak, cortico-cortical excitatory synapses governed by AMPAR-dynamics. Each network consists of 2,048 excitatory and 512 inhibitory leaky integrate-and-fire neurons fully connected through AMPAR-, NMDAR- and GABA_A_R-mediated synaptic transmission as in (Compte et al., 2000). Moreover, excitatory and inhibitory neurons were spatially distributed on a ring so that nearby neurons encoded nearby spatial locations. All connections were all-to-all and spatially tuned, so that nearby neurons with similar preferred directions had stronger than average connections, while distant neurons had weaker connections. Inhibitory-to-inhibitory and across-network connectivity was untuned. Intrinsic parameters for both cell types and all the connectivity parameters were taken from (Compte et al., 2000), except the following for networks holding up to two stimuli or capacity-2 networks (notation consistent with Compte et al., 2000):

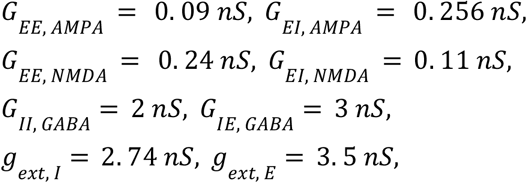

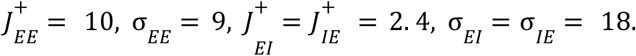

For networks holding up to 3 stimuli (capacity-3 networks),

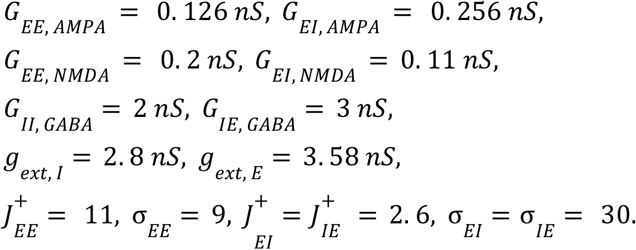

Connectivity across networks was determined by the following conductances (for unconnected simulations, these conductances were set to zero):

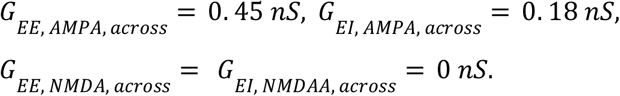

These parameters were adjusted to have within-network oscillations, which was accomplished by increasing the ratio between fast and slow excitation, supported respectively by AMPAR and NMDAR channels, as previously shown (Compte et al., 2000). The main dynamics described in this study were robust to a broad range of parameter values (Figure 1, 3, 4).

**Figure 1.**
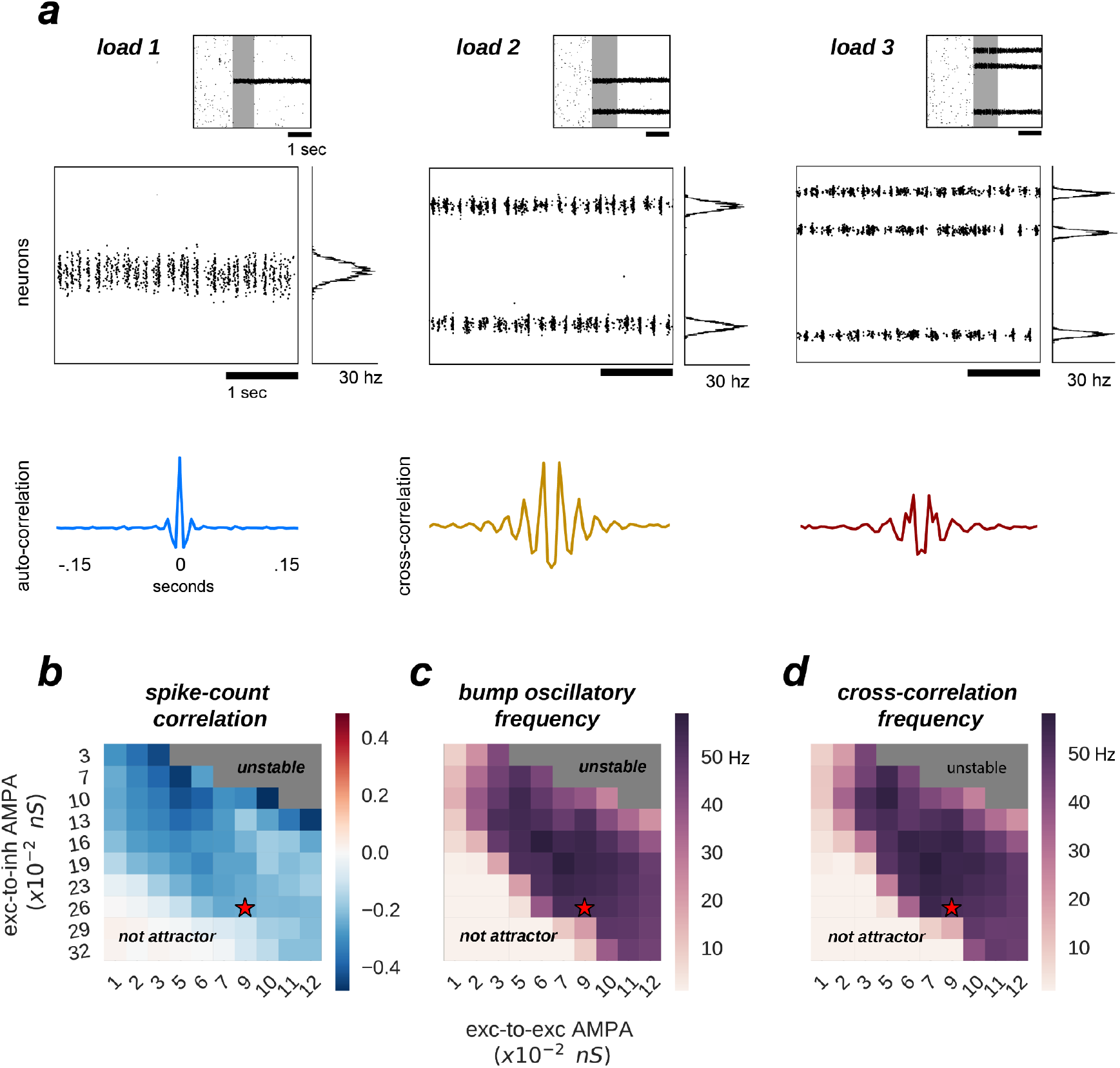
Multiple bumps are spontaneously anti-correlated. **a)** Raster plots of 3 sample simulations of load 1, 2 and 3 (top) and delay-period zooms (middle) show clear bump oscillatory activity, confirmed by correlation functions (bottom). Notably, irregular activity coexists with oscillatory dynamics. **b-d)** Anti-correlated oscillatory dynamics as excitation is manipulated in the network (AMPAR conductance for excitatory to inhibitory connections, y-axis, and for excitatory to excitatory connections, x-axis) in simulations with load 2 for the capacity-2 network used in Figures 3,4 (to facilitate comparisons). **b)** Anti-phase dynamics as measured by zero-lag cross-correlation between bumps. **c)** Dominant frequency of the auto-correlation function computed independently for each bump (computed with Fourier analysis). **d)** Dominant frequency of the cross-correlation between the two bumps. Red stars mark the parameter values of model simulations used throughout the study. Plots in b-d) summarize the dynamics of ∼10,000 simulations (total) of 100 different networks.

### Cross-correlations

For the cross-correlation analyses, we computed spike counts in bins of 5 ms, collapsing all neurons around the stimulus presentation location (here called a *bump*, ± 340 neurons). Moreover, we computed within- and across-network correlations by, respectively considering neurons in bumps from the same or different circuits. For the cross-frequency correlation plots (e.g. Fig 2b), we further computed the power spectrum of the resulting cross-correlation functions, averaged across all possible (only within- or only across-) pairs of bumps.

**Figure 2.**
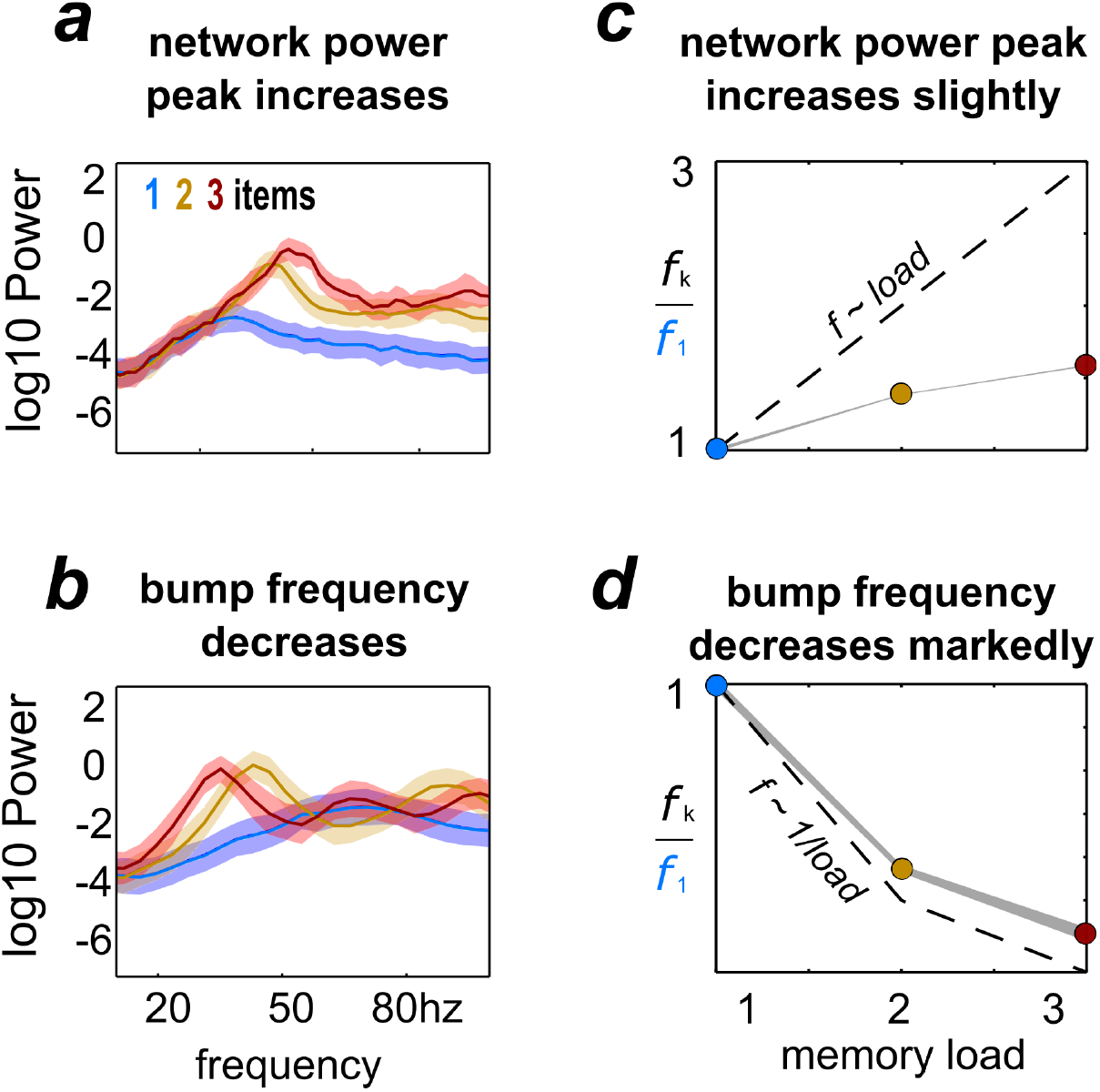
Load-modulation of network and bump oscillatory dynamics. Power spectrum of LFPs computed from simulations with increasing load (1-3), using the activity of the whole network **a)** or of each bump’s activity, **b). c, d)** Peak-frequency *f*_*k*_ computed from simulations with increasing load *k*, normalized to frequency *f*_1_from simulations with a single bump and computed from LFPs of the whole network activity (c) and from LFPs of each individual bump’s activity (d).

### Conversion of spikes into local field potentials

For the conversion of simulated spike trains into local field potentials, we convolved the aggregated spike times (*t*_*s*_) of all the neurons engaged in a bump (or in the network, depending on the analysis) with an alpha-function synaptic kernel:

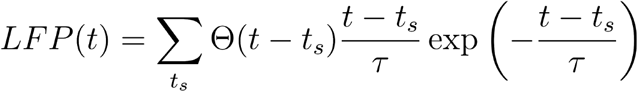

with Θ(*t*) being the Heaviside theta function, and τ = 5 *ms*.

### Phase-preservation index

To measure how an oscillating activity bump kept its oscillatory phase over multiple trials (*k*=1..*N*) of our simulation, we first converted spike times into local-field potentials (see above). Through wavelet analysis, we determined the phase *ϕ*^*k*^(*f*_0_, *t*) of the LFP at *f*_0_ = 30 Hz (the approximate frequency of oscillations in the network) at all time points *t* of the simulation, and then we used the phase-preservation index (PPI), a method originally developed by (Mazaheri and Jensen, 2006) for EEG data.

The PPI is defined by taking a reference time point (in our case *t*_*ref*_ = stimulus offset), and then computing the average consistency of the phases at the specific frequency of interest *f*_0_ with the rest of the time points:

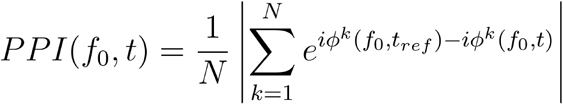

PPI values thus vary between 0 and 1, with 1 indicating perfect phase consistency.

## Results

### Working memory load modulates oscillation power and frequency

We built a computational network model of a local neocortical circuit, with excitatory and inhibitory spiking neurons *(leaky integrate-and-fire* neuron model) connected reciprocally via excitatory AMPAR-mediated and NMDAR-mediated synapses and inhibitory GABA_A_R-mediated synapses (see Methods). The ring-attractor network model was adjusted to support bump attractor dynamics with up to 3 simultaneous bumps (Edin et al., 2009), and further adjustment of the relative weights of AMPAR- and NMDAR-mediated currents was performed to set active reverberant neurons in the oscillatory regime (Wang, 1999). Using this computational model we started by investigating the dynamics that originated within each network.

In our model, multiple bumps showed anti-correlated oscillatory activity (Figure 1). As we stored more bumps in the network, lateral inhibition originating from simultaneous memories established anti-phase oscillatory dynamics during the memory period. These oscillatory dynamics were irregular, as illustrated in quickly dampened correlation functions (Figure 1a, bottom). Moreover, we found that the anti-phase behavior was robust in a wide range of values for AMPAR conductances (Figure 1b), consistently in the gamma range of frequencies (Figure 1c,d). Having seen these anti-phase dynamics between simultaneous bumps, we sought to contrast two opposite scenarios as we increased the number of stored memories (*memory load*). Under one alternative, bumps may oscillate at a fixed frequency irrespectively of load, so that the global network oscillation (adding up the activity of fixed-frequency out-of-phase bumps) would have a frequency that should increase linearly with memory load (scenario 1, dashed line Figure 2c). Alternatively, the network global oscillation could have a fixed frequency for different loads, and simultaneous bumps would take turns to fire in the available active periods. This would lead to halving each bump’s oscillation frequency as we double the memory load (scenario 2, dashed line in Figure 2d). We tested our model simulations to identify if our biophysical model adhered to one of these scenarios. To this end, we ran multiple simulations with three different loads (presenting 1, 2 and 3 separate bumps during the encoding cue period) and we computed power spectra from either the aggregate activity of the whole network (network power) or from separate populations centered around each presented target (bump power). We then extracted the frequency of the peak network and bump power to study their dependency with load. We found signatures of both scenarios (Figure 2a,b). As we increased the memory load, the overall network activity oscillated at slightly increasing frequencies (Figure 2a,c). In contrast, each bump, corresponding to different memories, oscillated at markedly slower frequencies as load increased (Figure 2b,d). We quantified which were the dominant dynamics by plotting both the network’s and each bump’s oscillating frequency against memory load. For better comparison, we normalized the frequency associated with different loads to the one of load 1. Moreover, we compared the effect of memory load against scenario 1 and 2 (dashed lines in Figure 2c,d). Qualitatively, we found that our network dynamics was more consistent with the latter.

We therefore conclude that our biophysical network maintains a relatively constant global oscillation as more items are loaded into memory, and individual memory oscillations instead start skipping cycles to sustain out-of-phase dynamics with other memories. Thus, the interplay between recurrent (fast) excitation and (slower) feedback inhibition acting locally is the basis of the oscillatory bump behavior. Moreover, we now show that anti-phase dynamics of simultaneous bumps occurs due to bump competition, accomplished by lateral inhibition. This competition increases with memory load, leading to longer periods of silence during the delay-activity of each bump. These dynamics generalize previous findings in simplified rate models (Pina et al., 2018), and extend them to biologically realistic ring attractor networks.

### Uniform coupling achieves feature binding

The binding between color and location is accomplished through the spontaneous synchronization of pairs of bumps across two networks connected via weak cortico-cortical excitation (Figure 3). In particular, we connected two ring-attractors in the regime described above with all-to-all, untuned excitatory connectivity. This connectivity was weak and it was mediated exclusively by AMPARs (Figure 3a), acting on all excitatory and inhibitory neurons. Interestingly, anti-phase dynamics within each network (as described above) was maintained robustly for a wide range of connectivity strength values (Figure 3e,f). Across networks, each bump’s activity was in phase with one bump in the other network (Figure 3b,c, black) but out of phase with the other (Figure 3b,c, red). On the majority of the simulations, this selective synchronization was maintained through the whole delay period (see Figure 3c,d for an example simulation). This dynamics is an interesting possible mechanism that binds and maintains the information of each presented stimulus. To this end, however, there are several aspects to resolve in relation to the encoding and decoding of this bound information.

**Figure 3.**
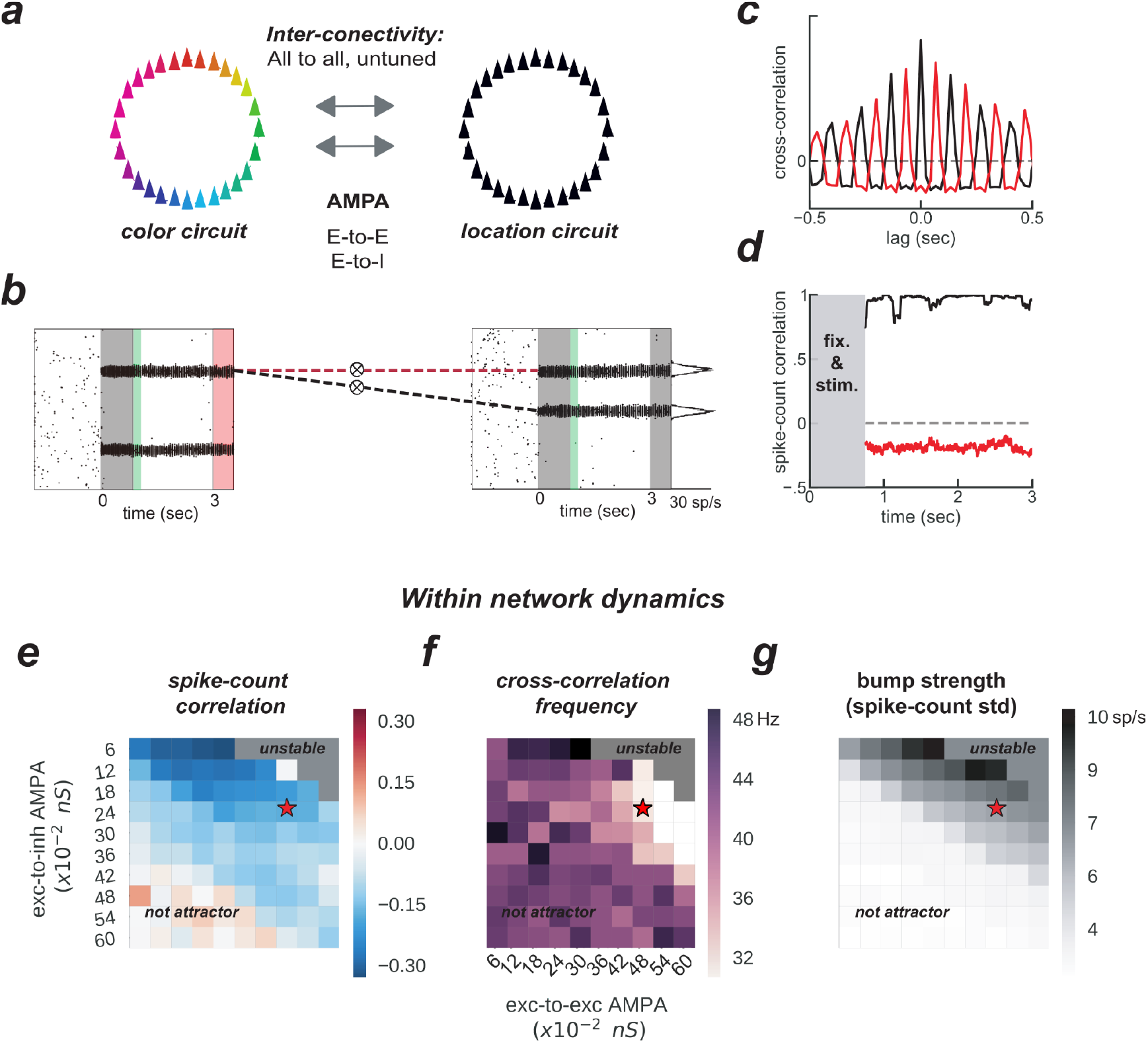
Feature-binding through weak, uniform coupling of 2 ring attractors. **a)** Schematics of the 2-network architecture, consisting of 2 ring-attractors with all-to-all, uniform connectivity. Each ring is able to store memories from one feature space (e.g. color or location) as activity bumps (Figure 1). **b)** One example simulation for the two networks. The pink-shaded area marks the period in which we read out the activity of the entire color network, while injecting current at one specific location in the location network (right gray-shaded area in the location rastergram, see main text for details about encoding/decoding). **c)** Cross-correlation computed between 2 pairs of bumps across networks (as marked with dashed red and black lines in panel b). Across networks, oscillating bumps synchronize in phase (black, positive zero-lag cross-correlation) or out of phase (red, negative zero-lag cross-correlation). **d)** Spike count correlation (in count bins of 5 ms and correlation windows of 100 ms) of both associations through the memory delay is stable for this simulation. **e)** and **f)** similar to Figure 1b,d, but manipulating AMPAR conductances across networks. e) Robustness of anti-phase dynamics within each network as measured by spike count correlation between bumps (Figure 1b). f) Dominant frequency of cross-correlation between the two bumps within each network (Figure 1d). **g)** Bump strength measured as standard deviation of spike-counts across model neurons at the end of the delay. e-g) summarize the dynamics of 22,000 simulations (total) of 100×2 networks. Stars indicate parameters and dynamical regime of network simulations shown in b-d.

On the one hand, synchronization selection was noise-induced in our simulations, resulting in across-networks associations between random pairs of bumps for different simulations. To control this association at the time of stimulus encoding, we stimulated strongly (7.5 times the intensity of sensory stimuli) and simultaneously 1 bump in each network for a brief period of 50ms (Figure 3b, and Figure 4a, green period), forcing these 2 bumps (1 in each network) to engage in correlated activity during the delay period. Nevertheless, this phase-locked dynamics could be broken by noisy fluctuations, leading to possible misbinding of memorized features and swap trials (Figure 4a,b).

**Figure 4.**
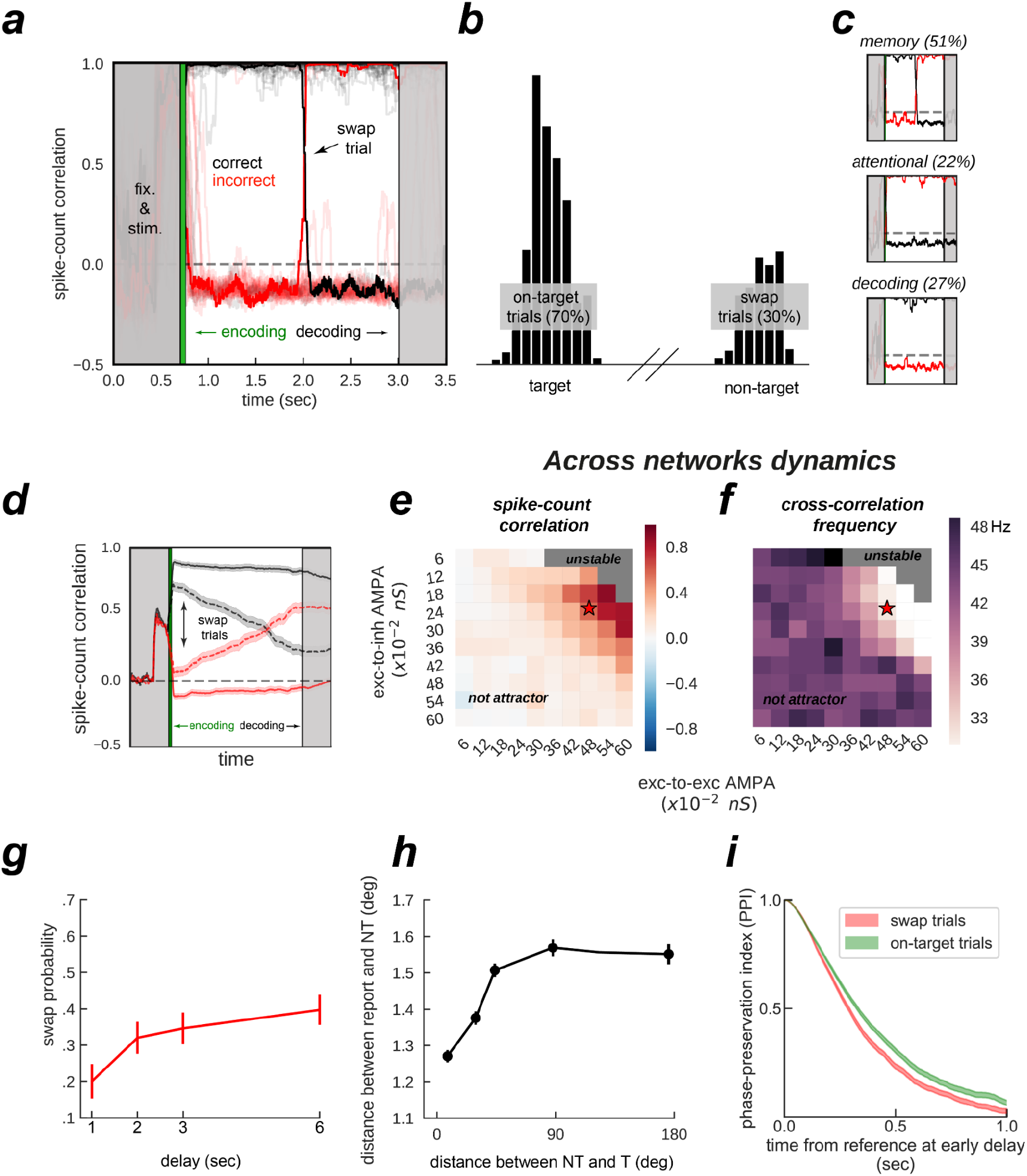
Encoding and decoding without temporal precision. **a)** Spike-count correlation (in count bins of 5 ms and correlation windows of 100 ms) during the delay for 20 sample simulations. During the encoding period (green), immediately after stimulus presentation, we bound two bumps, one from each network, by simultaneously stimulating them strongly. This ensured those two bumps were correlated through the trial more often than chance (black lines in the figures), and the other cross-network association synchronized mostly out-of-phase (red lines). On some trials (only one in a), noisy fluctuations reversed these correlations suddenly (swap trials). During the decoding period (light gray, on the right) we simulated the probe period of a working memory task, by stimulating the cued location (0.5 s) of one network, while decoding mean firing rates from the color network. **b)** Color readout histogram in 1,000 simulations. Bumps bound during encoding (target) were more likely to be read-out than unbound bumps (non-target). **c)** Three types of swaps: *memory swaps* (top), *attentional swaps* (middle) or *decoding swaps* (bottom). **d)** Same as a), averaging across all trials separately for swap and on-target trials, as defined by the decoder, shown in b). **e, f)** summary of the dynamics of 22,000 simulations (total) of 100 connected (x2) networks as a function of inter-network connectivity. **e)** Binding stability measured as the average spike-count correlation between initially bound bump pairs during the delay (black, in figures). **f)** Dominant frequency of the cross-correlation between bound bump pairs. Red stars mark the parameter values of the model used for sample simulations. **g)** Swap errors increase with delay duration and **h)** simulations where target (T) and non-target (NT) bumps are stored close-by increase swap errors, relative to when they are further apart. **i)** Swap-error trials (red, n=3,000), compared with on-target trials (green, n=3,000) in the model are associated with a lower phase consistency of oscillatory activity in the delay period, as measured with phase-preservation index (PPI, Methods) using early delay as the reference time point. Error-bars are bootstrapped standard errors (n=500).

On the other hand, our model raised the question of how this binding of information could reasonably be decoded without resorting to complex mechanisms for spike coincidence detection. In our task, the ‘behavioral’ output consisted in answering which ‘color’ was initially associated with a particular ‘location’, and this was accomplished by evaluating which bump of the color network maintained in-phase synchronization with the bump of the probed location at the end of the delay. We found that this did not require complex coincidence detection, but could instead be simulated in a rate formalism as follows. For each trial, we probed one *location* by stimulating weakly (¼ of stimulus intensity) corresponding neurons in the *location network* at the end of the delay. This simulated the visual presentation of a location probe at the end of the delay. This increased the firing rate of the corresponding location bump, and we found that it also resulted in an increase of activity of the associated, in-phase *color bump*. Finally, we extracted the behavioral output with a maximum likelihood decoder applied to the mean firing rate activity across the *color network* during the location-probing period (.5 s). Figure 4b shows color readouts from 1,000 of such simulated trials. Applying our encoding/decoding method to our simulations, resulted in 30% of trials wrongly associated with the non-target color (swap trials, Figure 4b). We then separated *swap* trials from *on-target* trials and computed the spike-count correlation in windows of 5 ms through the whole trial period (Figure 4d), and confirmed that on-target trials were in fact characterized by stable phase-locked activity, while the correlation between bumps in swap trials progressively approached the opposite dynamics (in-phase/anti-phase for the bound/unbound items, Figure 4d). Importantly, networks maintained synchronized in-phase dynamics for bound features robustly over a broad range of inter-network connectivity parameter values (Figure 4e,f). Additionally, we identified three sources of swap errors in our simulations, classified as *memory swaps* if the correct association based on in-phase bump synchronization changed abruptly during the delay (51% of the swap trials), *attentional swaps* if the wrong association was encoded during the encoding period (22%) or *decoding swaps* if the correct association was encoded and maintained during the memory period, but the decoding failed (27%). See Figure 4c for example simulations.

Together, our biologically-constrained simulations demonstrate that feature-binding can be robustly accomplished through selective synchronization. Crucially, encoding/decoding location-color associations was done without a *temporally precise code*, a long-standing limitation in the *binding by synchrony* framework (Shadlen and Movshon, 1999). Moreover, we identified 3 sources of swap errors. Based on these computational findings, we investigated model predictions that could be compared with existing data or could generate hypotheses for new experimental studies.

### Swap errors increase with delay and item competition

In our model, swap errors are induced by noisy fluctuations. This results in two behavioral predictions, congruent with previous findings (Pertzov et al., 2017; Schneegans and Bays, 2017; Emrich and Ferber, 2012). First, longer memory delays should increase the probability of a noisy fluctuation that is sufficiently large to induce a swap (Figure 4g). Second, Figure 4g shows how swap errors decrease with target to non-target distances. For very close locations, feedback inhibition is strongest, leading to “winner take all” dynamics between nearby bumps, explaining an increase of swap errors for such distances. For intermediate distances, similarly to (Almeida et al., 2015), simultaneous bumps interfere (repulsively and through their phase relationship, which is in this case less stable through the delay). Experimentally, these two regimes correspond to different scenarios. In the first case, one color is forgotten, while in the second scenario, there is an actual *swap* error. This prediction could be tested experimentally by probing the subject’s memory on all items, instead of just one (Adam et al., 2017).

In sum, our model is able to describe a previously found dependence of swap errors with delay duration and with target to non-target distance, and it offers mechanistic explanations for such dependencies.

### Neural prediction: swap trials show less phase preservation through the delay

Finally, abrupt changes in the phase relationship between oscillating bumps is the central mechanism of swap errors in our model (Figures 4a,b). Therefore, it is worth deriving a testable neurophysiological prediction from this mechanism. Additionally, because these changes in phase relationships are abrupt, they require experiments using techniques with high temporal resolution such as MEG or EEG. Intuitively, swap errors in our model simulations are characterized by inconsistent phase relationships between brain signals when comparing the beginning and the end of the delay period. We therefore considered applying an analysis that has been proposed to test phase consistency in EEG/MEG: the phase-preservation index (PPI, (Mazaheri and Jensen, 2006)). We first derived LFP signals from our network’s spiking activity (Methods). We then calculated the phase-preservation index (PPI, see (Mazaheri and Jensen, 2006) and Methods) at the end of the delay, relative to the beginning of the delay, and separately for on-target and swap trials defined “behaviorally” (Figure 4b). As we expected based on our model simulations (Figure 4), this analysis applied to our simulated data showed that trials containing swap errors had a lower PPI, compared to on-target trials (Figure 4g). This prediction can be tested with MEG/EEG data recorded from humans performing this task, based on an analysis of behavioral responses able to discriminate swap and correct error trials (Bays et al., 2009).

## Discussion

Aiming to account for swap-errors, a prominent source of multi-item working memory interference (Schneegans and Bays, 2019), we extended the ring-attractor model (Compte et al., 2000). Our biologically-constrained model offers a plausible mechanism for feature-binding. Briefly, the encoding and decoding of associations is accomplished through rate-coding, while their maintenance is accomplished through selective synchronization of oscillatory mnemonic activity. Oscillatory dynamics emerges naturally from bump competition, which increases with memory load and is in line with previous EEG experiments in humans (Roux et al., 2012) and LFP recordings from monkey PFC (Lundqvist et al., 2018). Finally, our model reveals different origins of swap errors (Mitchell et al., 2018; Pratte, 2019), how they depend on delay duration and inter-item distances (Pertzov et al., 2017; Schneegans and Bays, 2017; Emrich and Ferber, 2012), and predicts that phase-locked oscillatory activity during the memory periods should reflect swap errors.

### Other multi-area models for working memory

Our multi-area model adds to a large body of computational work (Min et al., 2020; Froudist-Walsh et al., 2020; Mejias and Wang, 2019; Engel and Wang, 2011; Ardid et al., 2007; Edin et al., 2009; Murray et al., 2017; Novikov et al., 2021; Ardid et al., 2010; Bouchacourt and Buschman, 2019) attempting to account for the distributed nature of working memory (Christophel et al., 2017). While several of these models have implemented across-area interactions through oscillatory dynamics (Ardid et al., 2010; Novikov et al., 2021), they did not attribute a clear mechanistic role to inter-area synchronization dynamics. This is in contrast to our model, where feature-binding in working memory is accomplished through selective synchronization of oscillatory activity in different brain areas.

### Comparison with previous binding models

Previously proposed models by (Pina et al., 2018) and (Raffone and Wolters, 2001) as well as our model are explicit implementations of the synchronization mechanism for feature binding in working memory. While similar in the approach, there are important differences. As argued by Schneegans and Bays (2019), a major difficulty with previous synchronization models was that they were unable to show their capacity of reproducing the rich phenomenology of working memory behavior that other models can explain. Our model, on the basis of its architecture with ring attractor models of spiking neural networks, overcomes the limitation of earlier discrete population models (Pina et al., 2018; Raffone and Wolters, 2001) and keeps all the demonstrated explanatory power that is characteristic of these attractor models, such as explaining several behavioral working memory biases in humans (Almeida et al., 2015; Nassar et al., 2018; Kilpatrick, 2018; Kiyonaga et al., 2017; Stein et al., 2020; Barbosa and Compte, 2018) and monkeys (Papadimitriou et al., 2015; Barbosa et al., 2020); as well as explaining key neurophysiological dynamics during working memory maintenance periods (see (Barbosa, 2017) for a short review) in humans (Edin et al., 2009; Kamiński et al., 2017) and monkeys (Wimmer et al., 2014; Sajad et al., 2016). Our model also goes beyond previous synchronization models in that (1) by virtue of its 2-ring architecture, it explicitly implements the storage of different features in independent systems or brain areas, as shown experimentally (Schneegans and Bays, 2019), and that (2) it provides a plausible rate-based readout mechanism of working memory associations without resorting to complex synchrony detection processes, a major difficulty for this sort of models (Shadlen and Movshon, 1999). Indeed, we show that our proposed mechanisms is robust to the noise inherent in spiking networks, which together with the need of precise spike coincidence detectors were major concerns of the binding through synchronized activity hypothesis in general (Shadlen and Movshon, 1999) and previous implementations in particular (Pina et al., 2018; Raffone and Wolters, 2001).

Thus, our model now brings back synchronization-based feature binding in working memory as a plausible alternative to recent conjunction binding proposals, such as the *binding pool* (Swan and Wyble, 2014) *and the conjunctive coding* model (Matthey et al., 2015; Schneegans and Bays, 2017). These models implement binding mechanisms that are fundamentally different from ours. In these models, binding of separated features is accomplished through conjunction neurons, which are neurons selective to mixtures of those features. While there is evidence for such neurons in the cortex (Fusi et al., 2016; Rigotti et al., 2013), their role in feature-binding is not clear, given the consistent evidence for separate feature storage underlying working memory binding (Delvenne and Bruyer, 2004; Olson and Jiang, 2002; Parra et al., 2011; Xu, 2002; Wheeler and Treisman, 2002; Fougnie and Alvarez, 2011; Bays et al., 2011b). Importantly, such a mechanism scales exponentially with the number of feature combinations, thus seemingly inconsistent with our ability to flexibly bind never seen combinations (Schneegans and Bays, 2019). However, it is to be noted that some conjunction models have mitigated this scaling problem through the construction of random conjunctions in an interposed network (Swan and Wyble, 2014; Bouchacourt and Buschman, 2019).

### Encoding with rate code

In our hybrid model, only the maintenance of associations is accomplished through correlated oscillatory activity or, in other words, relies on a *temporal code*. Instead, encoding and decoding of associations is achieved through a *rate code*. Encoding and decoding is accomplished by delivering flat pulses (i.e. without the need to be temporally precise) to both the to-be-bound features exclusively (*encoding)* or just to one of them (*decoding*).

Encoding the association between two different features through a pulse delivered simultaneously to each corresponding bump resembles the sequential encoding hypothesis in working memory (Wolfe, 1994; Bays et al., 2011a). Moreover, there is evidence that a mechanism combining sequential and parallel encoding is implemented in the brain when solving multi-item working memory tasks (Bays et al., 2011a). Our model implements such a combination. First, information about independent features arrives simultaneously to memory-encoding areas from upstream sensory areas. Then, the correct associations are sequentially encoded by brief excitatory pulses, possibly as a result of overt selective attention to each stimulus sequentially (Schoenfeld et al., 2014). Speaking to this, humans take longer to encode combined features than they take to encode the same amount of independent features (Schneegans and Bays, 2019).

### Decoding with rate code

Works modelling multi-item working memory though the storage of several bumps in a network (Krishnan et al., 2018; Wei et al., 2012; Nassar et al., 2018) - including our own (Almeida et al., 2015; Edin et al., 2009) - often used approaches that are biologically implausible to extract the location of one bump, while ignoring other simultaneously maintained bumps. Our approach, however, matches closely the “cueing” period of a multi-item working memory task, which consists of stimulating the “cued” locations while reading out from the whole color network population. Moreover, our encoding/decoding mechanism proposes that swap errors can be of different origins (attention, memory or decoding, Figure 4c). Indeed, experimental designs that require subjects to rate their confidence on a trial-by-trial basis show that swap errors occur both in high- and low-confidence trials, suggesting different origins (Mitchell et al., 2018; Pratte, 2019).

### Future work: towards biological plausibility of binding through dynamics

We found anti-phase dynamics within each network and phase-locking across networks, the central mechanisms for feature-binding in our model, to occur naturally in a broad range of parameters, indicating that the mechanisms proposed here do not require fine-tuning. Because our model is to some degree biologically constrained, it is a proof of concept that working memory binding through synchronized activity is *at least* possible to occur in the brain. In fact, we simulated noisy integrate-and-fire neurons, supporting that the central mechanism implemented in our model has some degree of robustness to noise.

Our model is, however, limited in several ways that could be addressed in future studies. First, we did not simulate trials demanding binding of load 3 or higher. We expect that the main challenges associated with that improvement will be the encoding of more associations. Second, we did not investigate how feature-binding is impacted by incoming distractors. Previous work has shown that oscillatory activity on different bands can play a role in filtering distractors (Dipoppa and Gutkin, 2013). Future work combining these models is necessary. Third, as a proof of concept, we only simulated 2 connected networks, while humans can encode and decode the association of many more features (Schneegans and Bays, 2019). Finally, the oscillatory regime in which our model is operating, in which neurons are strongly synchronized with the population rhythm (Figure 4c), however derived from biologically constrained neuronal models, is arguably not biological itself. While there is abundant evidence that neuronal populations show strong oscillatory dynamics in working memory (e.g. (Pesaran et al., 2002)), single neuron dynamics approaches a Poisson process (Softky and Koch, 1993; Compte et al., 2003) - therefore not oscillatory at this scale (but see (Lundqvist et al., 2016)). Early theoretical work (Brunel and Hakim, 1999; Brunel and Wang, 2003; Brunel, 2000) has demonstrated that such oscillatory dynamics at the population level can coexist with noisy, unsynchronized neurons when randomly connected. Future work that connects randomly connected networks that store multiple stable bump-attractors (Hansel and Mato, 2013), but operating in anti-correlated oscillatory activity such as in our simulations could be an appropriate avenue for the future work attempting to overcome these limitations.

## Data Availability Statement

The multi-area model as well as the code for measuring the phase-preservation index is available in https://github.com/comptelab/binding.

## Author Contributions

JB carried out the research. JB and AC conceived the research and wrote the manuscript. KS conceived the electrophysiological prediction. All authors edited and approved the final version of the manuscript.

## Funding

This work was funded by the Spanish Ministry of Science, Innovation and Universities and European Regional Development Fund (Refs: BFU2015-65315-R and RTI2018-094190-B-I00); by the Institute Carlos III, Spain (grant PIE 16/00014); by the Cellex Foundation; by the Generalitat de Catalunya (AGAUR 2014SGR1265, 2017SGR01565); and by the CERCA Programme/Generalitat de Catalunya. JB was supported by the Spanish Ministry of Economy and Competitiveness (FPI program) and by the Bial Foundation (Ref: 356/18). This work was developed at the building Centro Esther Koplowitz, Barcelona.

## Notes

### Competing Interest Statement

The authors have declared no competing interest.

